# Direct observation of interdependent and hierarchical kinetochore assembly on individual centromeres

**DOI:** 10.1101/2025.06.25.661565

**Authors:** Changkun Hu, Andrew R Popchock, Angelica Andrade Latino, Charles L Asbury, Sue Biggins

## Abstract

Kinetochores are megadalton protein machines that harness microtubules to segregate chromosomes during cell division. The kinetochores must assemble after DNA replication during every cell cycle onto specialized regions of chromosomes called centromeres, but the order and regulation of their assembly remains unclear due to the complexity of kinetochore composition and the difficulty resolving individual kinetochores *in vivo*. Here, by adapting a prior single-molecule method for monitoring kinetochore assembly in budding yeast lysates, we identify a sequential order of assembly and uncover previously unknown interdependencies between subcomplexes. We show that inner kinetochore assembly depends partly on outer kinetochore components, and that outer kinetochore branches do not assemble independently of one another. Notably, Mif2 assembly is a rate-limiting step that can be accelerated by binding to the Mtw1 subcomplex, thereby promoting rapid assembly of many inner and outer kinetochore components. The importance of controlling kinetochore assembly kinetics is supported by a Mif2 mutant lacking both autoinhibition and Mtw1 subcomplex binding activity, which leads to defective kinetochore-microtubule attachments when the centromeric histone variant Cse4 is overexpressed. Altogether, our work provides a direct view of kinetochore assembly and reveals highly interdependent regulatory events that control its order and timing.

**GRAPHICAL ABSTRACT:** 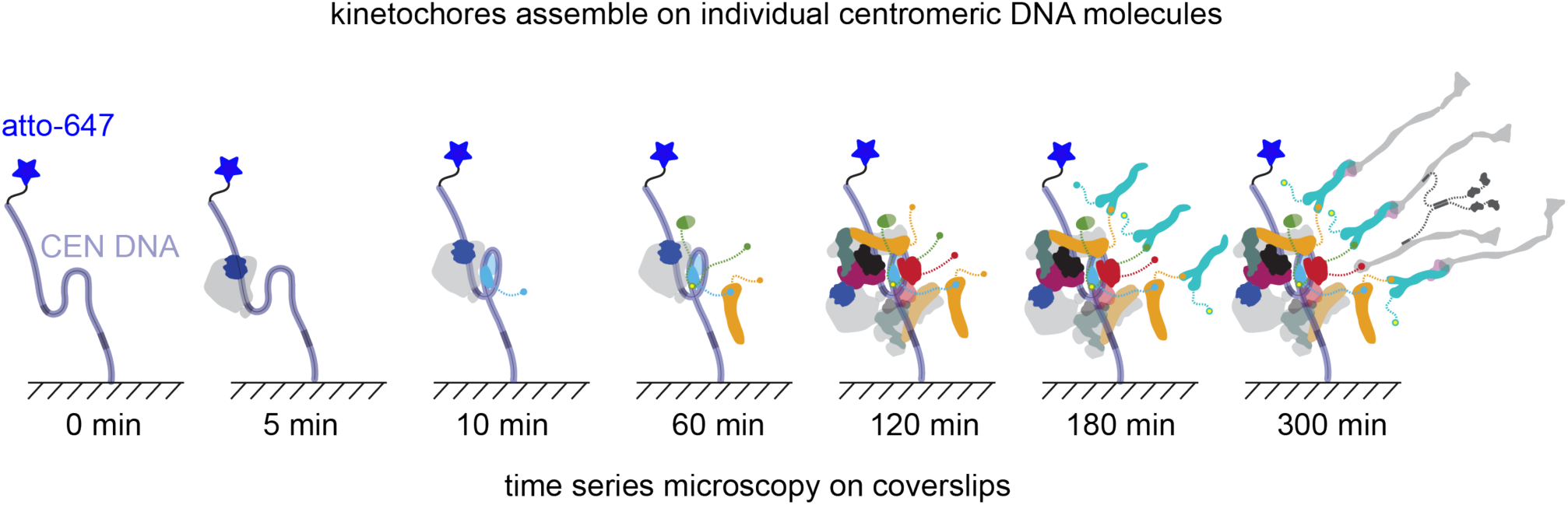

## Introduction

Organisms require accurate chromosome segregation to maintain their genomes and cellular fitness. Chromosome segregation is mediated by dynamic attachments between microtubules and kinetochores (1,2), protein machines that assemble at specialized sites on each chromosome called centromeres, whose locations are epigenetically determined by the presence of a histone H3 variant, commonly called CENP-A, and known as Cse4 in budding yeast (3,4). Kinetochores are dynamic, large assemblies composed of numerous unique proteins that exist in stable subcomplexes, each present in multiple copies per kinetochore (5–7). Inner kinetochore proteins assemble on the specialized centromeric chromatin structure and connect to the outer kinetochore proteins that mediate microtubule attachment.

A challenge in the field is to understand how kinetochores faithfully assemble specifically at the centromere every cell cycle. Budding yeast are an excellent system to study kinetochore assembly because they have small, sequence-specific centromeres that form a single centromeric nucleosome sufficient to mediate assembly of the entire kinetochore (8–12). Most budding yeast kinetochore subcomplexes are conserved (Table 1). A large complex called the <u>c</u>onstitutive <u>c</u>entromere <u>a</u>ssociated <u>n</u>etwork (CCAN) forms the inner kinetochore and bridges to the major microtubule binding activity that is provided in yeast by the Ndc80 and Dam1 subcomplexes (13,14). Recent structures of the budding yeast inner kinetochore show a highly interdigitated arrangement of CCAN subcomplexes surrounding the centromeric nucleosome (15,16) (Figure 1A). Disordered N-terminal portions of four inner kinetochore proteins, Cse4, Mif2, Cnn1, and Ame1, emanate from this structured core and form the bases of four biochemically distinct branches, which are thought to recruit outer subcomplexes independently from one another (17,18). Each branch potentially recruits the microtubule-binding Ndc80 subcomplex (Ndc80c), either directly, via Cnn1, or indirectly through the Mtw1 subcomplex (Mtw1c), via Cse4, Mif2 and Ame1. Because Cnn1 and Mtw1c both bind to the same site on Ndc80c (17,18), it has been assumed they act as independent receptors for Ndc80c in yeast (Figure 1A). The Ndc80c then recruits additional microtubule-binders, such as Stu2, and signaling components. This branched tree-like model is attractive because it explains (i) the hierarchy of colocalization dependencies observed *in vivo*, (ii) why biochemical purification of one subcomplex often copurifies others nearby in the hierarchy, and (iii) how a single centromeric nucleosome can recruit the large number of microtubule-binders required for persistent coupling to dynamic microtubule tips. However, direct studies of the kinetochore assembly process have been limited, so it is unclear whether the branches assemble independently. It is also unknown whether the kinetochore is built in a stepwise manner and what regulatory events are important for ensuring accurate assembly.

**Figure 1.**
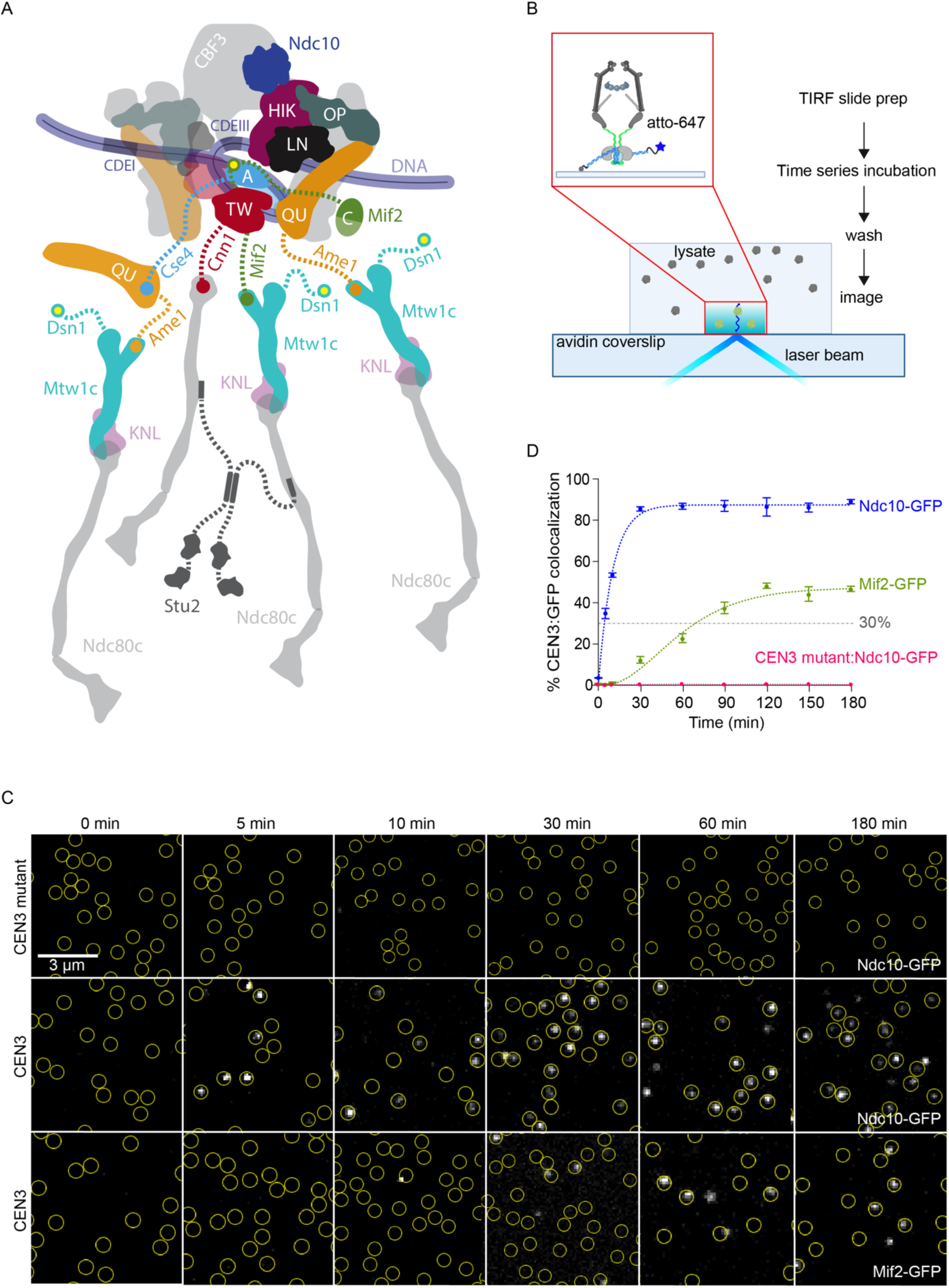
TIRFM assay reveals the assembly timing of kinetochore subcomplexes. (A) Schematic of the budding yeast kinetochore based on (16) demonstrating known interactions between inner kinetochore and outer kinetochore. Only proteins included in this study are labeled. Note that human homolog names are CENP-A, CENP-C, CENP-HIK, CENP-LN CENP-TW, CENP-OP, and CENP-QU (See Table 1). (B) Schematic describing time series TIRFM assay. Cells that contained fluorescent, GFP-tagged kinetochore proteins were treated with benomyl to depolymerize microtubules and arrest cells in mitosis. Lysates were prepared and then introduced into a flow-channel with fluorescent, Atto 647-tagged CEN3 DNA tethered sparsely onto the coverslip surface. After incubation for various times (0, 5, 10, 30, 60, 90, 120, 150, or 180 min), the lysates were washed out and then TIRFM was performed. (C) Representative TIRFM images of surface-tethered wild type or mutant CEN3 DNA molecules (at locations indicated by yellow circles) after incubation for the indicated times with lysates from strains carrying Ndc10-GFP (SBY22903) or Mif2-GFP (SBY22094). Ndc10-GFP accumulated rapidly and specifically on wild type CEN3 DNAs, whereas Mif2-GFP accumulated more slowly (GFP foci shown in grayscale). (D) Percentages of CEN3 or CEN3 mutant DNAs with colocalized Ndc10-GFP (SBY22903) or Mif2-GFP (SBY22094) versus time. Error bars represent the standard deviation over three biological repeats. At least 3,000 DNA molecules were imaged for each time point from each biological replicate. Fitting curves were generated using two-parameter kinetic equations as detailed in the methods.

**Table 1.**
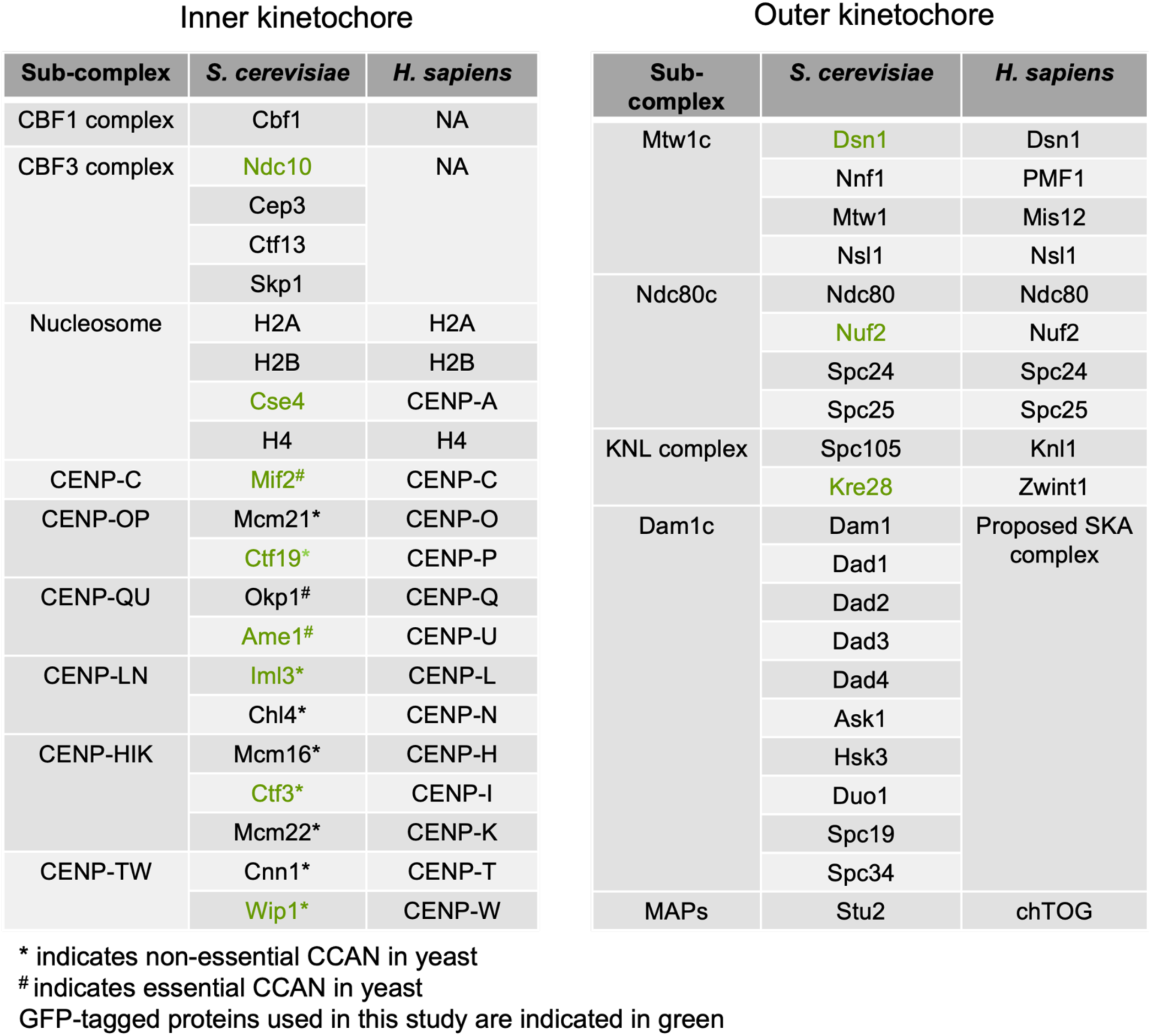
Kinetochore subcomplexes and proteins in *S.cerevisiae* and *H. sapiens*.

To investigate the underlying kinetics and temporal order of kinetochore assembly, we adapted a method to study yeast centromeric nucleosome assembly at the single molecule level using total internal reflection fluorescence microscopy (TIRFM) (19,20). With this method, we determined the relative order of assembly of a representative member of each kinetochore subcomplex. Our data reveal strong, unexpected interdependencies, such as an enhancement of CCAN assembly that depends on the interaction of Mif2 with Mtw1c. We also discovered that Mif2 autoinhibition by its N-terminus is limiting for kinetochore assembly and can be relieved by the binding of Mif2 to Mtw1c. Cells lacking the N-terminus of Mif2 are viable unless the centromeric histone is overexpressed, suggesting that the interdependencies we uncovered are particularly important when kinetochore assembly is challenged.

## Materials and methods

### Yeast strains and genetic modifications

The *S. cerevisiae* strains are described in Supplemental Table S1 and are derivatives of SBY3 (W303). Standard genetic crosses, media preparation, and microbial techniques were used. Cse4 was internally tagged with EGFP at residue 80, flanked by linkers (pSB1617), and expressed under its native promoter at the endogenous locus (SBY22195). Other kinetochore genes were modified to include endogenously tagged GFP (pSB3156) and auxin-inducible degrons (-IAA7) using standard PCR-based integration methods (Longtine et al., 1998), with modifications confirmed via PCR. The *mif2* mutants were generated by CRISPR-based integration using vectors carrying Cas9 (pSB3218) and the donor sequences *mif2-ΔN* (pSB3354) and *mif2^AME1N^* (pSB3237) (21). The primers and plasmids used to construct the strains are listed in Supplemental Table S2.

### Culture conditions

All liquid cultures were maintained in yeast peptone dextrose (YPD) medium. To synchronize cells in mitosis, 30 μg/mL benomyl was added to log-phase cultures for 2.5 hours, ensuring at least 90% of cells exhibited large-budded morphology. For strains with auxin-inducible degron (AID) alleles (*dsn1-AID*), cultures were treated with 500 μM indole-3-acetic acid (IAA) dissolved in DMSO with 30 μg/mL benomyl, following methods described (22–24). Strains with galactose-inducible alleles were initially cultured in 2% raffinose before being cultured in 2% galactose until they were harvested at the specified time points. To achieve G1 arrest, yeast cells were treated with 1 μg/mL alpha factor for three hours.

### Cell fitness serial dilution assay

Yeast strains were cultured overnight in YPD medium. Cell density was determined using a spectrophotometer (Bio-Rad) and adjusted to an OD_600_ of 1.0. Serial 1:5 dilutions were then prepared in water using a 96-well plate, and aliquots from each well were spotted onto YPD or YPD plates supplemented with benomyl (Sigma-Aldrich). The plates were incubated at the specified temperatures for two days before imaging.

### Cell extract

The procedure for preparing cell lysate for kinetochore assembly assays was previously described (24). Briefly, cells were cultured in liquid YPD medium until reaching logarithmic growth and arrested in mitosis with benomyl (30 μg/mL) in a total volume of 1 L, followed by harvesting via centrifugation at 4 °C. All subsequent steps were carried out on ice using buffers maintained at 4 °C. The cells were washed once with dH_2_O containing 0.2 mM PMSF and then with Buffer L (25 mM HEPES pH 7.6, 2 mM MgCl_2_, 0.1 mM EDTA pH 7.6, 0.5 mM EGTA pH 7.6, 0.1% NP-40, 175 mM K-Glutamate, and 15% Glycerol) supplemented with protease inhibitors (10 µg/mL each of leupeptin, pepstatin, and chymostatin, 0.2 mM PMSF) and 2 mM DTT. The cell pellets were snap-frozen in liquid nitrogen and lysed using a Freezer/Mill (SPEX SamplePrep). The lysis involved 10 cycles, each consisting of 2 minutes of grinding at 10 times per second, followed by a 2-minute cooling period. The resulting powder was weighed and resuspended in Buffer L using the formula: pellet weight (g) × 0.8 = volume (mL) of Buffer L. The resuspended lysate was thawed on ice, clarified by centrifugation at 16,100 × g for 30 minutes at 4 °C, and the protein-rich supernatant was extracted using a syringe. The lysate was then aliquoted and snap-frozen in liquid nitrogen. The resulting soluble whole-cell extracts typically had a protein concentration of 70–80 mg/mL. Pellets, powder, and lysates were stored at -80 °C.

### DNA template preparation

As previously detailed (22), plasmid pSB963 was utilized to produce the wild type CEN3 DNA templates, containing the centromere and surrounding pericentromeric sequences (277 bp total) flanked by additional linker DNA (212 bp upstream of CDEI and 302 bp downstream of CDEIII). Plasmid pSB972 was used to create the CEN3 mutant template for this study. PCR products were purified using the Qiagen PCR Purification Kit. For single-molecule TIRFM assays, the purified CEN DNA was diluted in TE buffer to a final concentration of approximately 1 ng/µL.

### TIRFM slide preparation

Coverslips and microscope slides were ultrasonically cleaned and PEG-passivated as described previously (19,20,25). Briefly, ultrasonically cleaned slides were treated with 3-Aminopropyl triethoxysilane, 99% (Sigma-Aldrich) before incubation with 1% (w/v) biotinylated PEG-SVA MW-5000K/mPEG-SVA MW-5000K (Laysan Bio) in flow chambers constructed using double-sided tape. Passivation was conducted overnight at 4 °C. Following passivation, the flow chambers were washed with Buffer L and then incubated with 0.3 M BSA/0.3 M Kappa Casein in Buffer L for 5 minutes. The chambers were subsequently washed with Buffer L, incubated with 0.3 M Avidin DN (Vector Laboratories) for 5 minutes, and washed again with Buffer L. Next, ∼100 pM CEN3 DNA template was added to the chambers for 5 minutes, followed by another wash with Buffer L.

For endpoint colocalization assays, slides were prepared as follows: flow chambers were filled with 100 µL of cell lysate containing the protein(s) of interest using pipetting and wicking with filter paper or Pasteur pipettes. After lysate addition, slides were incubated for 90 minutes at 25°C, then washed with Buffer L to remove the lysate. The chambers were subsequently filled with Buffer L containing an oxygen scavenger system (10 nM PCD, 2.5 mM PCA, and 1 mM Trolox) for imaging (26).

### TIRFM time series imaging and analysis

As previously detailed (19,20), all images were captured using a Nikon TE-2000 inverted RING-TIRF microscope equipped with a 100x oil immersion objective (Nikon Instruments) and an Andor iXon X3 DU-897 EMCCD camera. Images were acquired at a resolution of 512 x 512 pixels with a pixel size of 0.11 µm/pixel or 0.16 µm/pixel at a readout speed of 10 MHz. Atto-647-labeled CEN DNAs were excited at 640 nm for 300 ms, GFP-tagged proteins at 488 nm for 200 ms. Images were analyzed with the CellProfiler (4.2.6) to assess colocalization and quantify signals between the DNA channel (647 nm) and GFP channel (488 nm). Results were processed and visualized using FIJI (https://imagej.net/software/fij). Adjustments to example images, such as contrast or false color, were made using FIJI and uniformly applied across the entire field of view for each image. The lysates were washed off before imaging. At least 3,000 DNA molecules were imaged for each time point from each repeat. Error bars are standard deviation over three biological repeats.

Colocalization-vs-time data were fit (in Igor Pro 9) with one of the four equations given below, which represent simple kinetic reaction pathways having either one, two, three, or four irreversible transitions (irreversible steps). Such simple reaction pathways do not explicitly model the complexity of kinetochore assembly, but they closely approximate the kinetics of more complex reaction pathways, and they provide excellent fits to our data using only two adjustable parameters. For a simple one-step assembly pathway, with one irreversible binding event, the fraction of complete assemblies will vary across time according to equation:

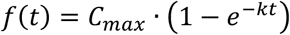

where *t* represents time (e.g., in min), 𝑘 represents the rate constant (min^-1^) for the transition, and *C*_*max*_ represents the maximum fraction achieved when *t* → ∞. For a simple two-step assembly pathway, with two irreversible transitions both having identical rate constants, 𝑘, the fraction of complete assemblies will vary according to equation:

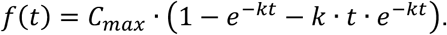

For a three-step assembly pathway, with three irreversible transitions all having identical rate constants, 𝑘, the fraction of complete assemblies will vary according to

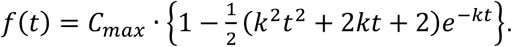

For a four-step assembly pathway, with four irreversible transitions all having identical rate constants, 𝑘, the fraction of complete assemblies will vary according to

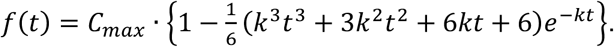

In general, the curve for a pathway with more step transitions exhibits a longer initial lag phase followed by a steeper rise to *C*_*max*_. We fit every dataset with all four equations, for one, two, three, and four steps, and then selected the best-fitting curve, with the minimum chi-squared deviation from the data. The rate constant, 𝑘, determines the time-scale over which 𝑓(*t*) rises to *C*_*max*_, but the best-fit values for 𝑘 are unimportant here because they do not correspond to any specific step in kinetochore assembly. Therefore, to compare curves we used 𝑇_30_, the time at which each curve reached a threshold of 30% (arbitrarily chosen). The best-fit values and uncertainties for *C*_*max*_ and 𝑇_30_, the numbers of step transitions in the best-fitting curves, and the resulting chi-squired deviations between the data and the fits are all included in Supplemental Table S3.

### In vivo microscopy and analysis

Fixed cell images were captured using a DeltaVision Ultra deconvolution high-resolution microscope (GE Healthcare) equipped with a 100X/1.42 PlanApo N oil immersion objective (Olympus) and a 16-bit sCMOS detector. Cells were imaged in Z-stacks through their entire volume, with 0.4 µm steps, using plane-polarized light, 488 nm and 568 nm illumination. All images were deconvolved using standard settings, and Z projections of the maximum signal from all channels were exported as TIFF files for analysis in FIJI.

Centromeric region puncta in cells were identified using 568 nm images of Ndc10-mCherry. Signal intensity within these regions was quantified for the 568 nm channel and the corresponding 488 nm channel on a per-cell basis. Total intensity in the 488 nm channel was normalized to the total 568 nm intensity for each cell, and the normalized values were averaged across approximately 33 cells per biological replicate, with three biological replicates analyzed per strain. Representative images shown from these experiments are maximum intensity projections across all Z images. Adjustments to example cell images, such as contrast, were made uniformly using FIJI.

### Immunoblotting

Cell lysates were made by beat-beat pulverization (Biospec Products) with glass beads in SDS buffer (27). SDS-PAGE gels were transferred to 0.45 µm nitrocellulose membrane using a wet transfer apparatus (Bio-Rad). The following commercial antibodies were used for immunoblotting: α-PGK1 (Invitrogen; 4592560; 1:10,000), α-Cse4 (9536L, 1: 500) (28). Secondary antibodies used were sheep anti-mouse antibody conjugated to HRP (GE Biosciences) at 1:10,000 or donkey anti-rabbit antibody conjugated to HRP (GE Biosciences) at 1:10,000. Antibodies were detected using SuperSignal West Dura Chemiluminescent Substrate (Thermo Fisher Scientific). Immunoblots were imaged with a ChemiDock MP system (Bio-Rad).

## Results

### A single molecule assay to monitor the kinetics of kinetochore assembly

We previously developed an approach using total internal reflection fluorescence microscopy (TIRFM) to observe the *de novo* assembly of individual centromeric nucleosomes and whole kinetochore particles in cell lysates (19,25). These efforts revealed molecular requirements for centromeric nucleosome assembly and directional asymmetry in the kinetochore-microtubule interface (19,20,25). However, the kinetics of kinetochore assembly beyond the centromeric nucleosome remained unexplored.

To examine kinetochore assembly kinetics more broadly, we adapted our TIRFM approach to monitor the recruitment of various kinetochore proteins onto individual centromeric DNAs via time-series imaging. Atto-647-labeled DNA containing the centromere and surrounding pericentromeric sequences from chromosome III (CEN3 DNA) as well as additional linker DNA (791 bp total) were sparsely tethered onto a coverslip surface to allow detection of individual CEN3 molecules (Figure 1B). One-step photobleaching confirmed that each CEN3 focus represented a single molecule (Supplemental Figure S1A-C) (29). To initiate kinetochore assembly on the surface-tethered CEN3 DNAs, we introduced yeast extracts containing GFP-tagged kinetochore proteins. We epitope tagged a representative component of each kinetochore subcomplex at its endogenous gene locus with GFP (Table 1, tagged proteins highlighted in green) and confirmed that the GFP tag did not affect cell fitness at various temperatures and in the presence of microtubule-destabilizing drug treatments (Supplemental Figure S1D). Because many CCAN proteins are non-essential, we used a more sensitive assay to determine whether the GFP tag affected their function. The non-essential CCAN genes become essential in a *dsn1-3A* mutant background (24,30), so we crossed the GFP-tagged genes to *dsn1-3A* to confirm they are functional (Supplemental Figure S1D). Once we had established a functional GFP tagged protein for each kinetochore subcomplex, we made lysates from cells arrested in mitosis with benomyl, to ensure the lysates were maximally competent for assembling kinetochores (24). To minimize possible variation in the rate of kinetochore assembly, we used a consistent lysate concentration (70 mg/mL) and incubation temperature range (21-23 °C) for the assay.

To establish the time series assay, we initially analyzed two inner kinetochore proteins (Ndc10-GFP and Mif2-GFP). Ndc10 is a component of the CBF3 complex that is required for assembly of the entire yeast kinetochore, including the Cse4 histone variant, suggesting that CBF3 binding is the initiating event (31–33). The Mif2 protein interacts directly with the centromeric nucleosome, suggesting it assembles after Cse4. We incubated each lysate with surface-tethered CEN3 DNAs for various durations (0, 5, 10, 30, 60, 90, 120, 150 and 180 min) and then washed the lysate away prior to imaging. Removing the lysate ensured stable protein binding and a high signal-to-noise ratio, free from background due to out-of-focus GFP or autofluorescence. Multicolor TIRFM images were then recorded, and colocalization single-molecule spectroscopy (CoSMoS) was performed (Figure 1C) (19,25,34,35). The fraction of wild type CEN3 DNAs decorated with GFP increased as the incubation time was extended, until reaching a maximum. The rates of increase and the maximum levels of colocalization were markedly different for the two kinetochore proteins (Figure 1C-D and Supplemental Figure 1E). Colocalization of Ndc10-GFP on wild type CEN3 DNAs rose above 50% in just 10 min and reached a maximum near 90% in 30 min. In contrast, Mif2-GFP assembled much more slowly than Ndc10-GFP, requiring 30 min to achieve 10% colocalization and 120 min to reach a maximum around 45%, consistent with the known requirement for Ndc10 to localize Mif2 (31–33). As a negative control, we also tested a mutant CEN3 DNA containing a 3-bp substitution that abolishes kinetochore assembly (24,36–38). The fraction of CEN3 mutant DNA molecules decorated with Ndc10-GFP never exceeded 1%, confirming the specificity of Ndc10 binding.

We fit the colocalization versus time data for each protein with kinetic curves representing simple reaction schemes (as detailed in Materials and Methods). To compare the assembly kinetics between proteins, we used 𝑇_30_, the time at which each curve reached a threshold of 30%, and the maximum colocalization, *C*_*max*_, which occurred when the curve plateaued at a steady value. The Ndc10-GFP colocalization plateaued at *C*_*max*_ = 87 ± 1% and reached 30% colocalization at 𝑇_30_ = 4.0 ± 0.2 min, consistent with the essential, early role Ndc10 plays in initiating kinetochore assembly (32). The Mif2-GFP data plateaued at *C*_*max*_ = 47 ± 1% and reached 30% colocalization at 𝑇_30_ = 70 ± 4 min, indicating that Mif2 assembles later than Ndc10, after a significant delay (Figure 1D). Altogether, these data show that the assembly of GFP-tagged kinetochore components specifically onto individual CEN3 DNAs can be monitored in time-series TIRFM, and that the rates and efficiencies of assembly are different for various components.

### Inner-kinetochore subcomplex incorporation is consistent with hierarchical assembly

Prior work indicates that kinetochore proteins are often expressed as biochemically stable, obligate heteromeric subcomplexes, most with two to four protein members (5,6,39). Obligate members of the same subcomplex are expected to assemble onto the kinetochore simultaneously; distinct (non-obligate) subcomplexes, however, are thought to assemble hierarchically, with DNA-contacting inner subcomplexes assembling first and outer subcomplexes incorporating later, only after their direct contacts in the inner kinetochore have already assembled (39). Co-localization dependencies and biochemical co-immunoprecipitations are consistent with this view, but the relative timing of subcomplex incorporation has not been measured directly, owing to the high-speed and complexity of the assembly process and the difficulty of distinguishing single kinetochores *in vivo*. We therefore extended our approach to compare assembly kinetics for representative members of every core kinetochore subcomplex, beginning with Cse4 (centromeric nucleosome) and the other five subcomplexes that make up the CCAN (Table 1).

The second component to assemble after Ndc10 was the centromeric histone Cse4 that reached 30% at 𝑇_30_ = 12 ± 1 min, only 7 minutes after Ndc10, and reached a maximum of *C*_*max*_ = 71 ± 1% by 90 min (Figure 2A). Next was the essential CCAN subcomplex, CENP-QU, represented by Ame1, reaching a similar maximum (*C*_*max*_ = 79 ± 3%) but more slowly, with 𝑇_30_ = 33 ± 3 min. The remaining four non-essential CCAN subcomplexes, assayed using GFP-tagged Ctf19 (CENP-OP) (36), Ctf3 (CENP-HIK) (40), Iml3 (CENP-LN) (41) and Wip1 (CENP-TW) (17,39), all assembled later and with lower maximum efficiency, following kinetic curves roughly similar to each other and to that of Mif2, with *C*_*max*_ = 38 ±1% (Ctf19), 56 ± 1% (Ctf3), 37 ± 1% (Iml3), and 49 ± 2% (Wip1) and 𝑇_30_ = 74 ± 10 min (Ctf19), 68 ± 2 min (Ctf3), 72 ± 3 min (Iml3), and 89 ± 7 min (Wip1), respectively (Supplemental Table S3). The similar timing of the non-essential CCAN components suggests they may be pre-assembled. We attempted to test more directly for the arrival of pre-assembled CCAN components using two-color, fast time-lapse imaging, in lysates where two different proteins were labeled with different fluorophores. Unfortunately, the relative fluorophore photostability and autofluorescence from the lysate precluded reliable detection of co-arrivals when obligate members of the same subcomplex were tagged (e.g., Okp1-GFP and Ame1-mKate).

**Figure 2.**
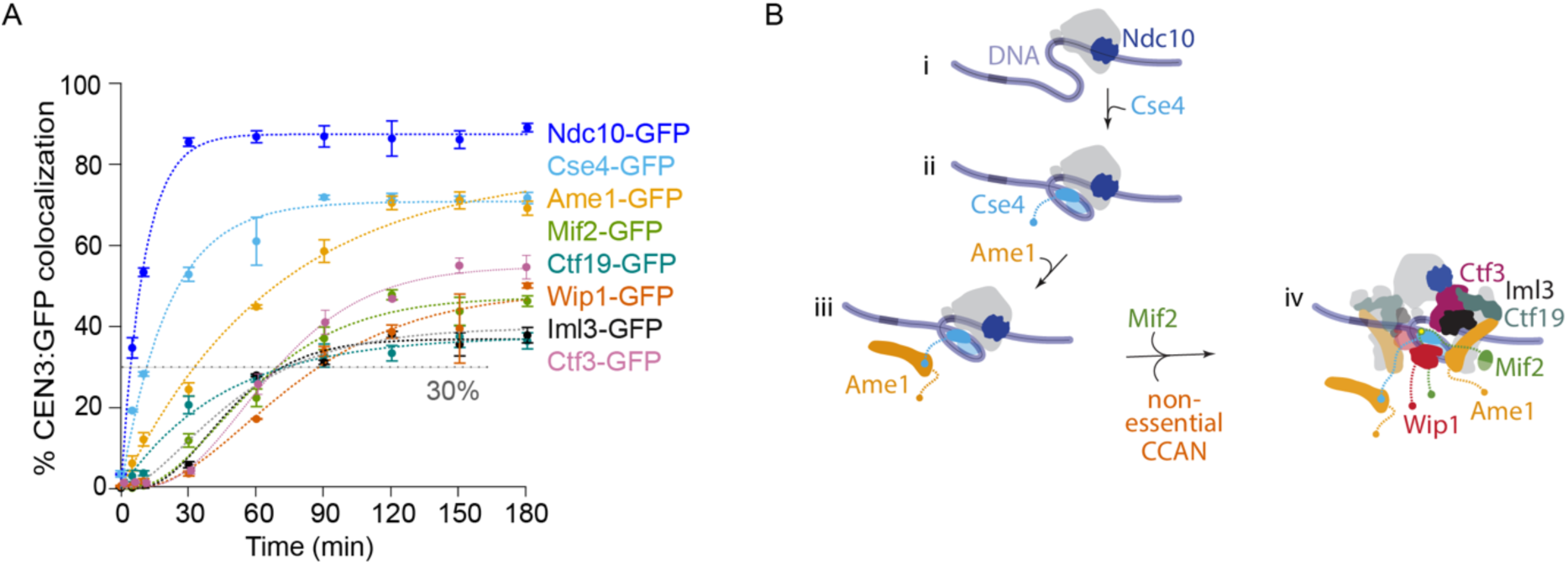
Centromeric nucleosome assembly is followed by incorporation of the CCAN subcomplexes. (A) Percentages of CEN3 DNAs with colocalized GFP-tagged Ndc10 (SBY22903), Cse4 (SBY22195), Ame1 (SBY22119), Mif2 (SBY22094), Ctf19 (SBY22116), Wip1 (SBY22207), Iml3 (SBY22199), or Ctf3 (SBY22203). Ndc10-GFP and Mif2-GFP data are replotted from Figure 1C for comparison. Error bars represent the standard deviation over three biological repeats. At least 3,000 DNA molecules were imaged for each time point from each biological replicate. Control experiments using the assembly-blocking CEN3 mutant DNA confirmed that all of these GFP-tagged proteins assembled specifically on wild type CEN3 DNA (Supplemental Figure S2). (B) Schematic model illustrating the assembly order of the inner kinetochore. (i) Ndc10 binds to CEN3 DNA to initiate kinetochore assembly. (ii) The Cse4 centromeric nucleosome forms. (iii) Ame1 binds to the centromeric nucleosome. (iv) Mif2 and the non-essential CCAN proteins assemble.

Taken together, these results are consistent with a hierarchical inner-kinetochore assembly process, initiated by Ndc10 and followed by Cse4 histone incorporation. One of the essential CCAN subcomplexes, represented by Ame1, then binds to the centromere followed by the other essential CCAN protein, Mif2, and the remaining four non-essential CCAN subcomplexes (Figure 2B). In general, the *C*_*max*_ tended to decrease the further a subcomplex was from the CEN3, consistent with our prior observations of de novo kinetochore assembly in lysates assayed by immunoblotting, where centromere-distal components assembled with reduced efficiency compared to centromere-adjacent components (24).

### Relieving autoinhibition of the Mtw1 subcomplex improves outer kinetochore assembly

We next sought to analyze outer kinetochore assembly using a representative GFP-tagged protein from three outer kinetochore subcomplexes, Dsn1 (Mtw1c) (28,39,42), Nuf2 (Ndc80c) (43,44), Kre28 (KNL subcomplex) (42), as well as the Stu2 protein (human chTOG homolog) (23,31). Unfortunately, the overall colocalization for the outer kinetochore subcomplexes was too low (∼2 to 15%) for reliable kinetic analyses (Figure 3A). The inefficient assembly of the outer kinetochore is consistent with our previous study where de novo assembly was measured in a bulk assay by immunoblotting (24). To overcome this limitation, we added phosphomimetic aspartic acid substitutions at two serine residues in the Dsn1 protein, S240D, S250D (*dsn1-2D*), which promotes outer kinetochore assembly by relieving autoinhibition of the Mtw1 subcomplex to promote its binding to Mif2 (Figure 3B) (24,30,45). Consistent with bulk kinetochore assembly assays (24), lysates from *dsn1-2D* strains produced much higher maximum colocalizations for GFP-tagged outer kinetochore proteins in the time-series TIRFM assay with *C*_*max*_ = 54 ± 2% (Dsn1-2D), 57 ± 1% (Nuf2), 52 ± 2% (Kre28), and 48 ± 1% (Stu2) (Figure 3C-D). Dsn1-2D assembly reached 30% at 𝑇_30_ = 56 ± 5 min (Figure 3D). Nuf2 assembled more slowly than Dsn1-2D, reaching 30% at 𝑇_30_ = 88 ± 4 min (Nuf2). Finally, Kre28 and Stu2 assembled last with 𝑇_30_ = 146 ± 9 min (Kre28) and 134 ± 6 min (Stu2), respectively. Altogether, these data confirm that relieving autoinhibition of the Mtw1 subcomplex by a phosphomimetic mutation significantly improves outer kinetochore assembly in the assay, and they suggest that the Mtw1 subcomplex likely assembles first, followed by the Ndc80 subcomplex, and then the KNL subcomplex and the Stu2 protein.

**Figure 3.**
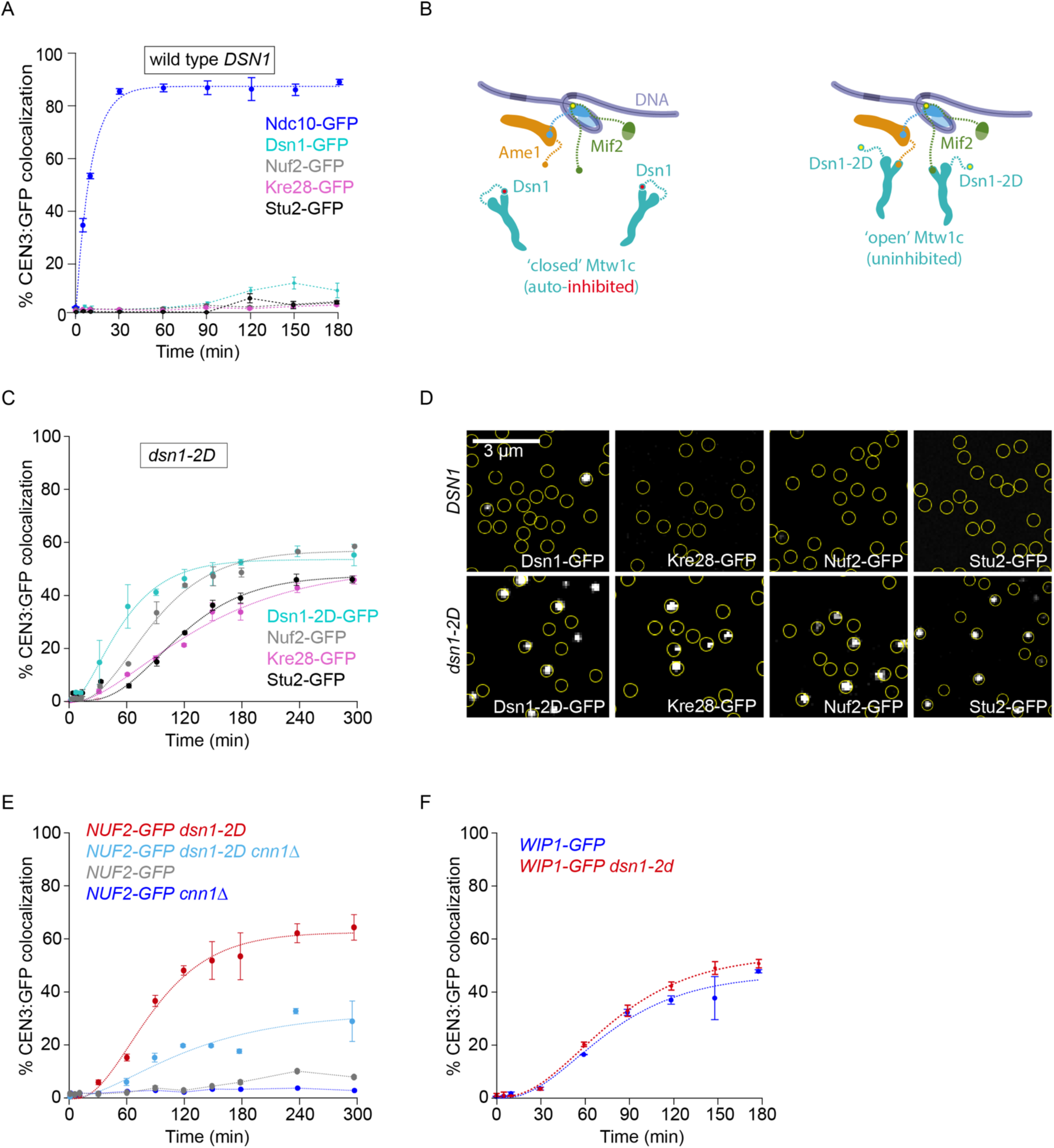
Relieving autoinhibition of the Mtw1 subcomplex with *dsn1-2D* improves outer kinetochore assembly. (A) Percentages of CEN3 DNAs with colocalized Ndc10-GFP (SBY22903) and GFP-tagged outer kinetochore proteins Dsn1 (SBY22153), Nuf2 (SBY23256), Kre28 (SBY24188) and Stu2 (SBY22135). Ndc10-GFP data are replotted from Figures 1C and 2A for comparison. (B) Schematic illustrating the regulation of the Mtw1 subcomplex by Aurora B phosphorylation. Mtw1 binding to Mif2 is autoinhibited (“closed”, left panel) until Dsn1 is phosphorylated by the Aurora B kinase (“open”, right panel). Once open, Mtw1 can bind to the N-terminus of Mif2. (C) Percentages of CEN3 DNAs with colocalized outer kinetochore proteins Dsn1-2D-GFP (SBY22159), Nuf2-GFP (SBY23258), and Kre28-GFP (SBY24190) in the *dsn1-2D* background. (D) Representative TIRFM images of GFP-tagged outer kinetochore proteins (grayscale) after 180 min incubation of CEN3 DNAs (yellow circles) with indicated lysates. Dsn1-GFP, Nuf2-GFP, Kre28-GFP, and Stu2-GFP are shown in the *DSN1* background (top; using lysates from strains SBY22153, SBY23256, SBY24188, and SBY22135, respectively) or in the *dsn1-2D* background (bottom; using strains SBY22159, SBY23258, SBY24190, SBY22133, respectively). (E) Percentages of CEN3 DNAs with colocalized Nuf2-GFP in *dsn1-2D* (SBY23258, data replotted from Figure 3D), *dsn1-2D cnn1*Δ (SBY24414), *DSN1* (SBY23256, same data from 3A), and *DSN1 cnn1*Δ (SBY24412) lysates. (F) Percentage of CEN3 DNA with colocalized Wip1-GFP in *DSN1* (SBY22207, data replotted from Figure 2A) and *dsn1-2D* (SBY22205) strains. All error bars represent the standard deviation over three biological repeats. At least 3,000 DNA molecules were imaged for each time point from each biological replicate.

### Assembly of Ndc80c onto the Cnn1 and Mtw1 subcomplexes is highly interdependent

The Ndc80c has two distinct kinetochore receptors in yeast, the Mtw1 and Cnn1 subcomplexes, that are thought to compete for binding to Ndc80c (18,46). Although Mtw1c did not assemble well in wild type lysates (less than 15% colocalization for Dsn1 after 300-minute incubation), Wip1 (representing the Cnn1/Wip1 yeast homologue of the CENP-TW complex) assembled relatively efficiently (*C*_*max*_ = 49 ± 2%). It was therefore surprising that Ndc80c did not assemble well (∼8% for Nuf2), suggesting that most of the centromere-bound Cnn1 in wild-type lysates did not recruit Ndc80c. To directly test the contribution of Cnn1 to Ndc80c recruitment, we measured Nuf2 assembly in lysates from mutant *cnn1Δ* cells. Nuf2 assembly dropped from ∼8% in wild type to ∼3% in the mutant *cnn1Δ* lysates (Figure 3E), confirming that the Cnn1 pathway in wild type lysates is only partially active. In *dsn1-2D* lysates, Nuf2 assembled to much higher levels, *C*_*max*_ = 57 ± 1%, consistent with the higher levels of assembled Mtw1 subcomplex. Strikingly, when we analyzed the contribution of Cnn1 to Ndc80c recruitment in *dsn1-2D* lysates, we found that Nuf2 recruitment dropped by half, to *C*_*max*_ = 31 ± 2%, in *cnn1Δ dsn1-2D* lysates (from 57% to 31%). The levels of Wip1 were similar in *DSN1* and *dsn1-2D* lysates (*C*_*max*_= 49 ± 2% versus 59.5 ± 0.8%, respectively, Figure 3F), indicating that neither Cnn1 nor the Mtw1 subcomplex were sufficient on their own for efficient recruitment of Ndc80c. Instead, maximal recruitment of Ndc80c relied on a strong, unexpected interdependency between both the Cnn1 and Mtw1 receptors. The *dsn1-2D* modification, which in a simple independent-assembly scenario would solely improve recruitment by the modified Mtw1 subcomplex, also caused a large increase in recruitment of Ndc80c by Cnn1.

### Relieving Mtw1 autoinhibition promotes CCAN and Mif2 assembly

In the simplest models of hierarchical assembly, stable incorporation of inner DNA-proximal subcomplexes would occur independently of the outer subcomplexes, and altering the outer Mtw1 subcomplex (e.g., using *dsn1-2D*) would not be expected to affect the assembly of inner subcomplexes. To test this hypothesis, we measured assembly kinetics in *dsn1-2D* lysates for the same inner-kinetochore subcomplexes that we had measured earlier in *DSN1* cells. As expected, the kinetics for GFP-tagged Ndc10 and Cse4 were similar to those measured in wild type lysates (*C*_*max*_ = 88 ± 1% and 76 ± 1%, respectively; and 𝑇_30_ = 2.6 ± 0.2 min, 12 ± 1 min, respectively) (Figure 4A-B). However, Ame1 showed slightly lower colocalization and slower assembly kinetics the in *dsn1-2D* background (*C*_*max*_ = 77 ± 3% and 𝑇_30_ =49 ± 4 min) for reasons that are unclear (Figure 4C). The GFP-tagged non-essential CCAN components, Ctf19, Iml3, and Ctf3 assembled in *dsn1-2D* lysates with similar timing relative to wild type lysates (𝑇_30_ = 53 ± 8, 52 ± 2, and 69 ± 23 min). However, they reached higher maximum efficiencies (*C*_*max*_ = 80 ± 5%, 60 ± 1%, and 69 ± 1%) (Figure 4D-F and Figure 3F). The protein that exhibited the biggest change was Mif2, which assembled much earlier in *dsn1-2D* lysates (𝑇_30_ = 26 ±1 min), now preceding Ame1 instead of following it and reaching higher maximum colocalization than in wild type lysates (*C*_*max*_ = 70 ± 1%) (Figure 4G-H and Figure 5A). These observations show that the timing and efficiency of inner subcomplex assembly can be strongly affected by modifications of outer subcomplexes, further indicating that kinetochore assembly occurs through a network of interdependent steps, rather than a simple sequence of independent events.

**Figure 4.**
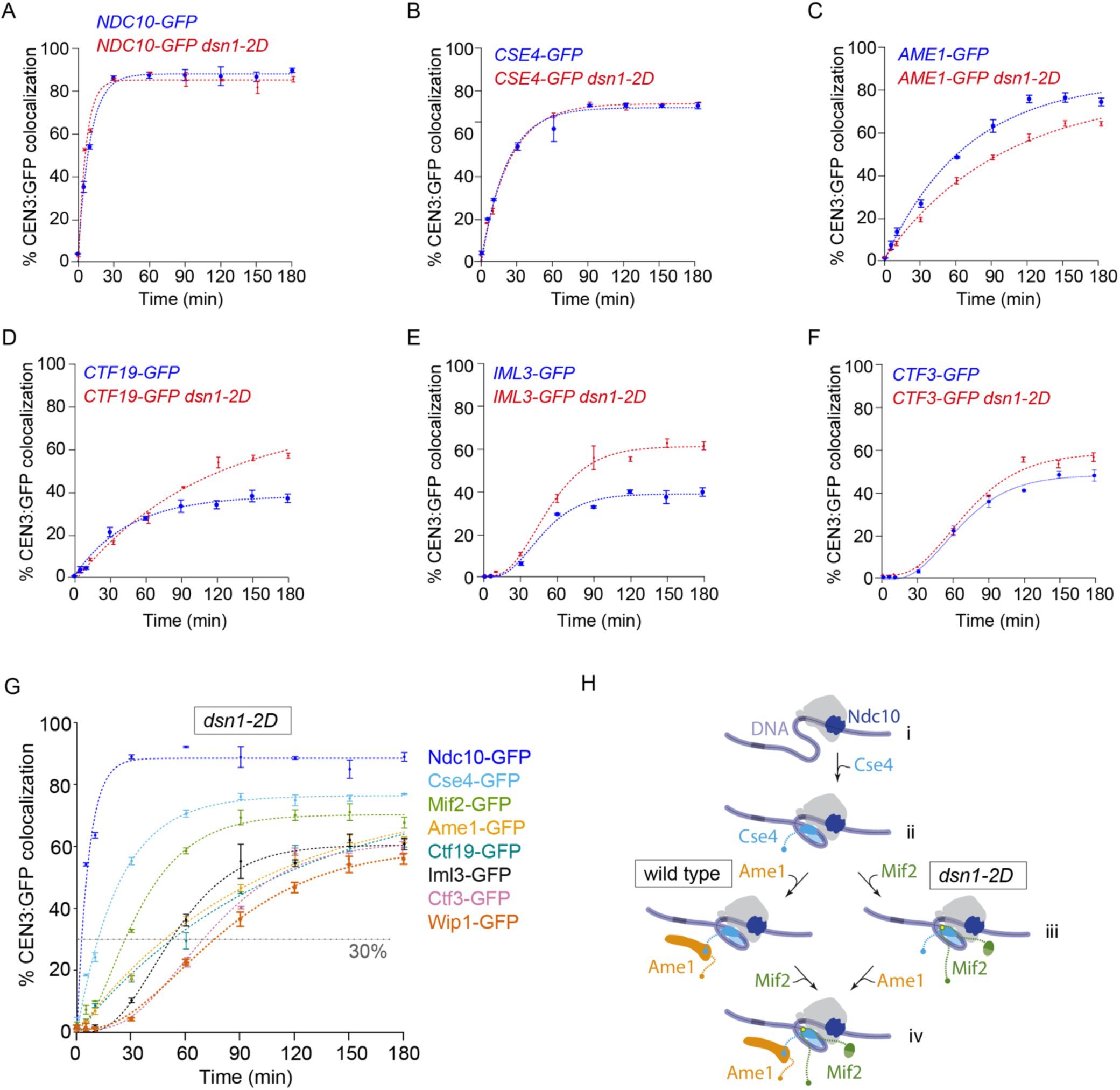
Relieving autoinhibition of the Mtw1 subcomplex with *dsn1-2D* improves assembly of CCAN components. (A-F) Percentages of CEN3 DNAs with colocalized GFP-tagged inner kinetochore proteins after assembly in DSN1 and dsn1-2D lysates. All DSN1 data are replotted from Figure 2A for comparison. GFP-tagged proteins in the dsn1-2D background include Ndc10 (SBY22905), Cse4 (SBY22193), Ame1 (SBY22117), Ctf19 (SBY22114), Iml3 (SBY22197), and Ctf3 (SBY22201). (A) Percentages of CEN3 DNAs with colocalized GFP tagged inner kinetochore proteins after assembly in dsn1-2D lysates, including the same data from Figure 4A-F, plus data for Mif2-GFP (SBY22092) and Wip1 (SBY22205). (B) Schematic model illustrating the change in assembly order of the inner kinetochore in dsn1-2D lysates (right pathway) compared to wild type (left pathway, from Figure 2B). (i) Ndc10 binds to CEN3 DNA to initiate kinetochore assembly. (ii) The Cse4 centromeric nucleosome forms. (iii) Mif2 binds to the centromeric nucleosome. (iv) Ame1 and the rest of non-essential CCAN proteins (not included) assemble. All error bars represent the standard deviation over three biological repeats. At least 3,000 DNA molecules were imaged for each time point from each biological replicate.

**Figure 5.**
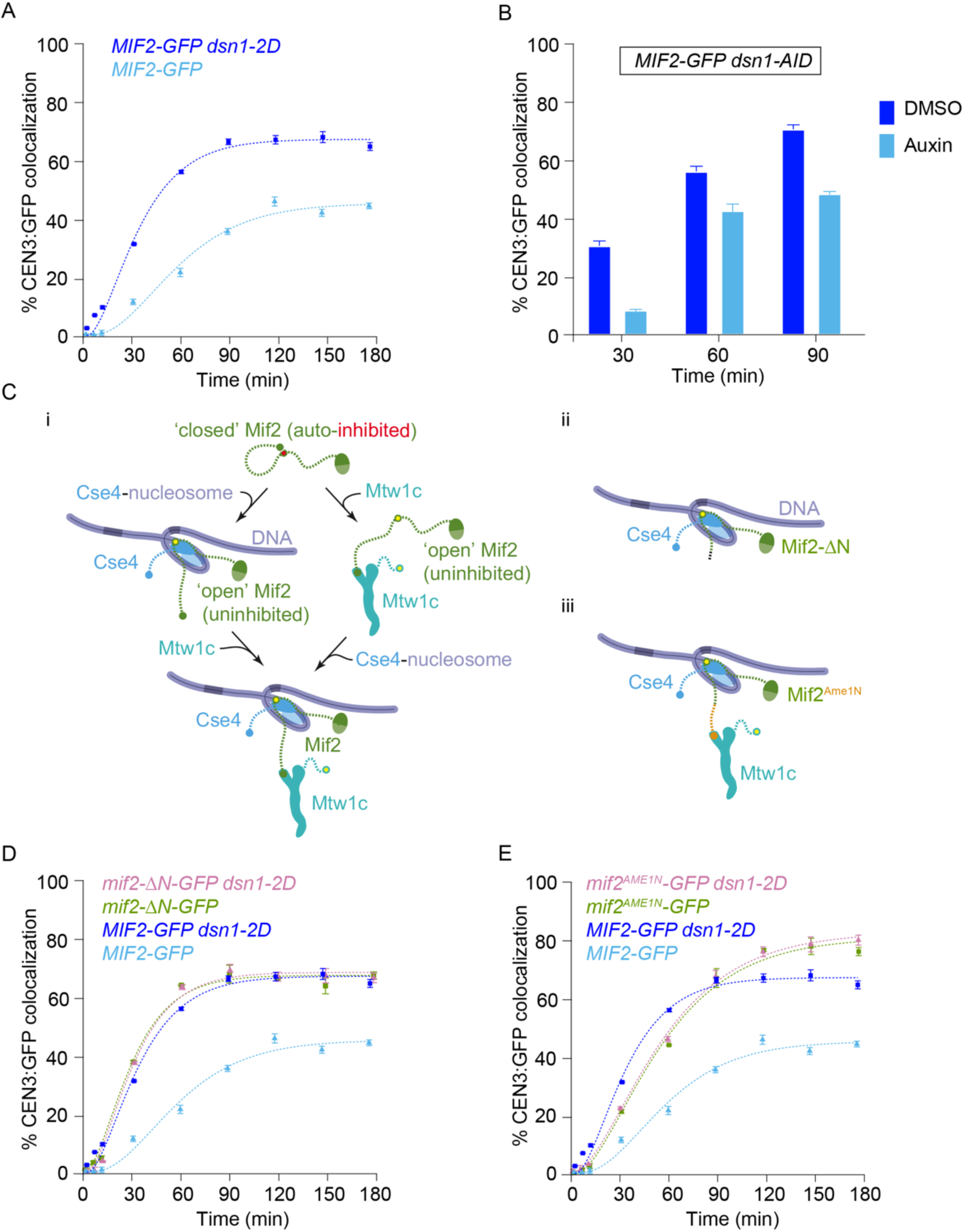
Relieving autoinhibition of the Mtw1 subcomplex with *dsn1-2D* also alleviates Mif2 autoinhibition. (A) Percentages of CEN3 DNAs with colocalized Mif2-GFP after assembly in *DSN1* (SBY22094, data replotted from Figure 2A) or *dsn1-2D* lysates (SBY22092, data replotted from Figure 4G). (B) Percentages of CEN3 DNAs with colocalized Mif2-GFP after assembly in lysates made from a *dsn1-AID* strain (SBY22872) treated with auxin to deplete the Dsn1-AID protein, or with DMSO as a control. (C) Schematics illustrating Mif2 autoinhibition. (i) Mif2 autoinhibition (closed) is relieved by binding to Cse4 (left pathway), making Mif2-N available (open) for binding to Mtw1c. Mif2 autoinhibition can also be relieved by binding to Mtw1c (right pathway), making Mif2 available for Cse4 binding (open). (ii and iii) Deleting the N-terminus of Mif2 (Mif2-ΔN, ii) or replacing it with an N-terminal fragment of Ame1 (Mif2^Ame1N^, iii) can relieve Mif2 autoinhibition. When the Ame1 N-terminus is grafted onto Mif2, it also restores binding to Mtw1c. (D) Percentages of CEN3 DNAs with colocalized wild type Mif2-GFP or colocalized truncation mutant Mif2-ΔN-GFP after assembly in either *DSN1* or *dsn1-2D* lysates (from strains SBY22094, SBY23248, SBY22092, and SBY23250). Data from strains with wild type Mif2-GFP are replotted from Figures 1C and 4G. (E) Percentages of CEN3 DNA with colocalized wild type Mif2-GFP or colocalized swap mutant Mif2^Ame1N^-GFP after assembly in either *DSN1* or *dsn1-2D* lysates (from strains SBY22094, SBY23252, SBY22092, and SBY23254). Data from strains with wild type Mif2-GFP are replotted from Figures 1C and 4G. All error bars represent the standard deviation over three biological repeats. At least 3,000 DNA molecules were imaged for each time point from each biological replicate.

### Relieving Mtw1c autoinhibition via *dsn1-2D* also relieves Mif2 autoinhibition

Adding phosphomimetic *dsn1-2D* substitutions to the outer Mtw1 subcomplex improved assembly of the inner Mif2 protein, so we tested whether the improved assembly required Mtw1c. To do this, we depleted Mtw1c using a *dsn1-AID* strain. For reasons that are unclear, Mif2 assembly in the *dsn1-AID* lysates that were not treated with auxin was higher than the wild-type strain. However, auxin treatment led to a reduction in Mif2 incorporation, especially at earlier time points (Figure 5B). Together, these data confirm that Mif2 assembly kinetics are regulated by Mtw1c.

A previous study showed that, like Mtw1c, Mif2 is also autoinhibited (Figure 5C-i) (21). The Mif2 N-terminal region, which directly recruits Mtw1c, can instead form autoinhibitory, intramolecular bonds to other regions of Mif2 that normally anchor it to the centromeric nucleosome. When Mif2 interacts with Cse4 nucleosomes, its autoinhibition is relieved, allowing its N-terminal region to bind Mtw1c (21). Our discovery that *dsn1-2D* improves Mif2 assembly suggested that Mtw1c binding may also help stabilize Mif2’s "open" status, promoting its binding to Cse4 (see right side of Figure 5C-i.) To further test this idea, we analyzed two Mif2 mutants lacking autoinhibition, Mif2-ΔN and Mif2^Ame1N^ (Figures 5C-ii and 5C-iii). We first deleted residues 2 through 35 from the N-terminus of Mif2, creating Mif2-ΔN, and thereby eliminating its ability to bind Mtw1c as well as its autoinhibition (21,47). Compared to wild type, the GFP-tagged truncation mutant Mif2-ΔN assembled onto centromeric DNAs much more quickly (𝑇_30_ = 38 ± 1 min for truncated Mif2-ΔN versus 70 ± 1 min for wild type) and to a higher final colocalization percentage (*C*_*max*_= 84 ± 1% for truncated Mif2-ΔN, versus 47 ± 1% for wild type) (Figure 5D), consistent with autoinhibition in the wild type Mif2 normally limiting its assembly at the centromere. Moreover, assembly kinetics for the Mif2-ΔN truncation mutant were indistinguishable regardless of whether the lysate contained wild type *DSN1* or phosphomimetic *dsn1-2D*. These observations indicate that the truncation mutant Mif2-ΔN is already uninhibited and therefore does not require the relief of autoinhibition that wild type Mif2 normally receives by associating with Mtw1c.

We next tested another Mif2 mutant, Mif2^Ame1N^, that lacks autoinhibition but retains its interaction with Mtw1c (21). The Mtw1 subcomplex interacts with the N-terminus of Ame1 in a manner that is not expected to be strongly regulated by *dsn1-2D* (45), so we made a Mif2 swap mutant where the N-terminus of Mif2 was replaced with the N-terminus of Ame1 (creating Mif2^Ame1N^; Figure 5C-iii) (21). Similar to Mif2-ΔN, the Mif2^Ame1N^ swap mutant assembled significantly earlier than wild-type Mif2 (𝑇_30_ = 25 ± 1 min for swap mutant Mif2^Ame1N^ versus 70 ± 4 min for wild type) (Figure 5E). Assembly kinetics for the swap mutant Mif2^Ame1N^ were indistinguishable regardless of whether the lysate contained wild type *DSN1* or phosphomimetic *dsn1-2D*, consistent with the Ame1 interaction with Mtw1c being independent of regulation by *dsn1-2D* (Figure 5E). Together, these data indicate that Mtw1c binding relieves the autoinhibition of wild type Mif2, allowing it to assemble more quickly and efficiently onto the centromere.

### Interactions between Mif2 and Mtw1c promote CCAN incorporation

We next asked whether the accelerated arrival of Mif2 affects the assembly of other CCAN proteins. To do this, we analyzed assembly kinetics for GFP-tagged CCAN components in lysates carrying the *mif2-ΔN* truncation mutant, which lacks the N-terminus and cannot bind to Mtw1c. Ctf19 assembly was normal (*C*_*max*_ = 40 ± 2%, 𝑇_30_ = 94 ± 15 min), but other non-essential CCAN components showed lower efficiency in *mif2-ΔN* lysates (*C*_*max*_= 29 ± 1% for Iml3, 41 ± 1% for Ctf3, and 22 ± 1% Wip1), and slower kinetics (𝑇_30_ = 174 ± 33 min for Iml3, 107 ± 3 min for Ctf3, and >180 min for Wip1, which never reached 30%) relative to lysates with wild type Mif2 (Figure 6A-D). These data indicated that the N-terminus of Mif2 enhances CCAN assembly. To test whether this enhancement was due to Mtw1 binding to Mif2, we used the Mif2^Ame1N^ swap mutant. In lysates carrying the Mif2^Ame1N^ swap mutant, the Ctf19 subcomplex again assembled normally (*C*_*max*_= 43 ± 3%, 𝑇_30_ = 84 ± 17 min), but the other non-essential CCAN proteins, Iml3, Ctf3 and Wip1, now showed higher colocalization efficiency (*C*_*max*_= 68 ± 1% for Iml3, 80 ± 1% for Ctf3, and 67 ± 1% for Wip1) compared to lysates with wild type Mif2. Two CCAN proteins, Iml3 and Wip1, also exhibited faster kinetics (𝑇_30_= 51 ± 2 min for Iml3 and 60 ±1 min for Wip1) (Figure 6A-D). These observations indicate that interactions between Mtw1c and Mif2 promote assembly of many components of the inner kinetochore, a result that is inconsistent with the simplest models, where inner components would assemble completely independently of outer components.

**Figure 6.**
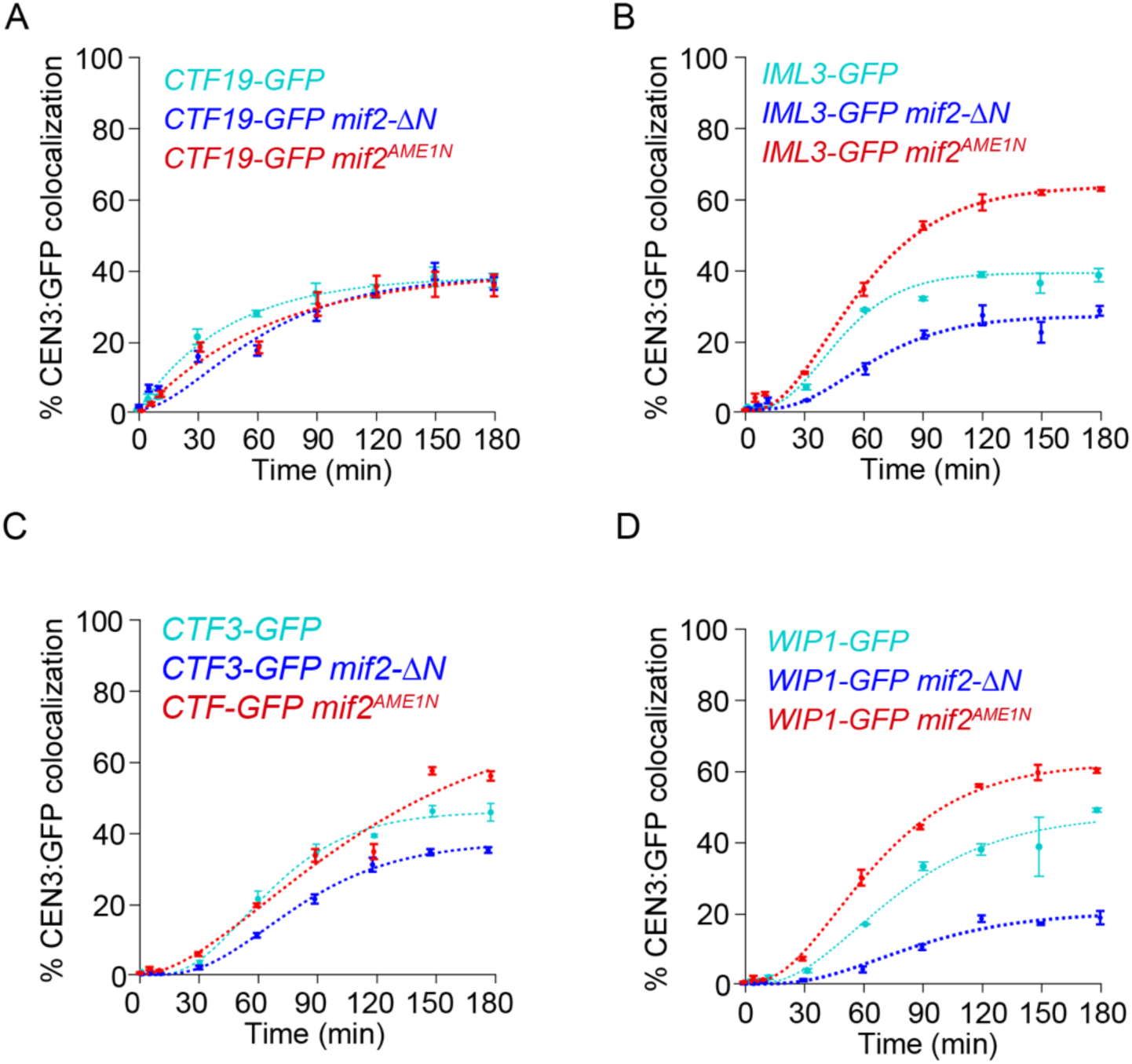
Interactions between Mif2 and the Mtw1 subcomplex improve assembly of non-essential CCAN components. (A) Percentages of CEN3 DNAs with colocalized Ctf19-GFP in wild type (SBY22116), *mif2-*Δ*N* (SBY24438), and *mif2^AME1N^* (SBY24420) lysates. (B) Percentages of CEN3 DNAs with colocalized Iml3-GFP in wild type (SBY22199), *mif2-*Δ*N* (SBY24436), and *mif2^AME1N^* (SBY24418) lysates. (C) Percentages of CEN3 DNAs with colocalized Ctf3-GFP in wild type (SBY22203), *mif2-*Δ*N* (SBY24426), and *mif2^AME1N^* (SBY24422) lysates. (D) Percentage of CEN3 DNA with colocalized Wip1-GFP in wild type (SBY22207), *mif2-*Δ*N* (SBY24430), and *mif2^AME1N^* (SBY24424) lysates. All data from strains with wild type Mif2 are replotted from Figure 2A for comparison. Error bars represent the standard deviation over three biological repeats. At least 3,000 DNA molecules were imaged for each time point from each biological replicate.

### The N-terminus of Mif2 becomes important in vivo when Cse4 is overexpressed

Although the Mif2 mutants lacking autoinhibition are viable, it was previously shown that overexpression of the Mif2^Ame1N^ mutant led to chromosome missegregation (21). We wondered whether the early arrival of Mif2 might also become critical in cells where Cse4 is overexpressed because it could promote ectopic kinetochore formation and/or titrate proteins from the endogenous kinetochores. To test this, we overexpressed Cse4 in *MIF2-GFP* and *mif2-ΔN-GFP* strains using a galactose-inducible promoter. Cells were plated on glucose and galactose media and, consistent with previous studies, Cse4 overexpression by itself did not alter cell fitness (21,48,49). However, there was a strong growth defect when Cse4 was overexpressed in *mif2-ΔN* mutant cells, consistent with Mif2 autoinhibition becoming important when the histone variant is misregulated (Figure 7A and Supplemental Figure S3). To determine whether Mif2 localization was altered, we analyzed GFP-tagged Mif2 or Mif2-ΔN in cells containing Ndc10-mCherry to mark the endogenous kinetochores. Cells were released from G1 into galactose media to induce Cse4 overexpression and we performed microscopy on mitotic cells. In wild type large budded cells, we nearly always observed two kinetochore foci, consistent with yeast kinetochores clustering into two foci when kinetochores biorient and come under tension (50). However, when Cse4 was overexpressed in *MIF2-ΔN-GFP* cells, we often observed more than two Mif2-GFP and Ndc10-mCherry foci (Figure 7B-C). The Mif2-GFP and Mif2-ΔN-GFP foci were colocalized with Ndc10-mCherry in greater than 99% of the cells, although the mCherry intensity was much lower than the GFP signal. Because Ndc10 specifically binds to CDEIII, the additional foci appear to represent endogenous, not ectopic kinetochores, and we hypothesized that they might represent impaired attachments between kinetochores and microtubules. Unattached kinetochores trigger the spindle assembly checkpoint (51–53), so we asked whether the *mif2-ΔN* cells overexpressing Cse4 activate the checkpoint. We released cells from G1 in the presence of galactose to induce Cse4 overexpression and analyzed the percentage of cells that entered anaphase 120 and 150 min after release. By 150 min, more than 60% of *MIF2-GFP*, *mif2-ΔN-GFP,* and *MIF2-GFP pGAL-CSE4* cells had entered anaphase. In contrast, less than 20% of the *pGAL-CSE4 mif2-ΔN* cells had progressed into anaphase. To determine whether this slow progression was due to spindle assembly checkpoint activation, we deleted the *MAD3* checkpoint gene. Deletion of *MAD3* rescued anaphase entry, indicating that the cells overexpressing Cse4 with Mif2-ΔN triggered the checkpoint, strongly suggesting they have defective kinetochore-microtubule attachments (Figure 7D). Given that we did not find evidence for ectopic kinetochores, it is likely that Mif2-ΔN overexpression affects endogenous kinetochore assembly. Regardless of the mechanism, these data suggest that the N-terminus of Mif2 becomes important for kinetochore function when there is excess Cse4.

**Figure 7.**
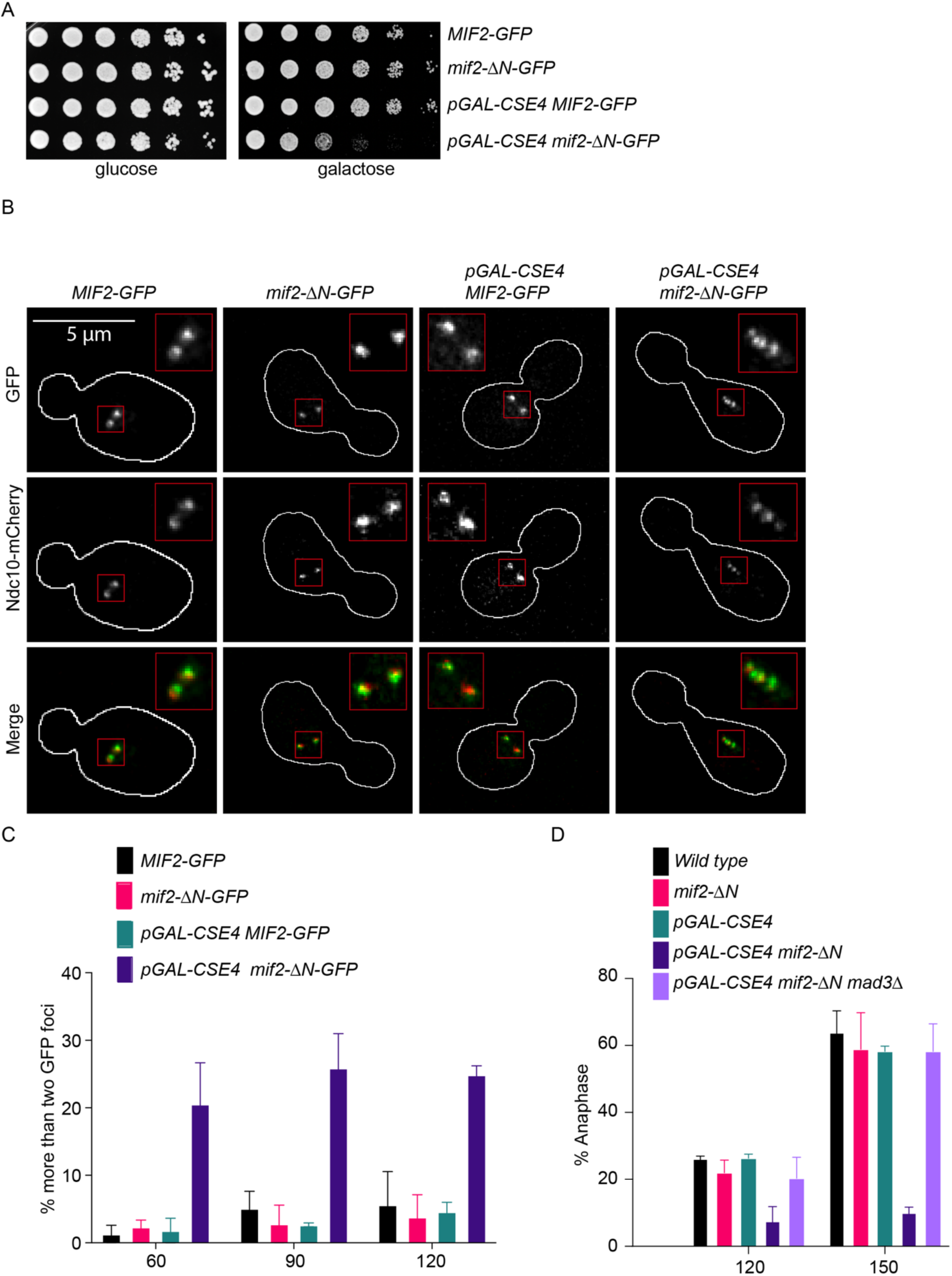
Cse4 overexpression in *mif2-*Δ*N* cells generates defective kinetochore-microtubule attachments. (A) Five-fold serial dilutions of *MIF2-GFP* (SBY22095), *mif2-*Δ*N* (SBY23249), *pGAL-CSE4 MIF2-GFP* (SBY24220)*, and pGAL-CSE4 mif2*-Δ*N-GFP* (SBY24222) yeast strains. Cells were grown on glucose and galactose plates and incubated at 23 °C for 48 hours. (B) Representative images of Mif2-GFP, Mif2-ΔN-GFP, and Ndc10-mCherry in wild type (SBY24202*), mif2-*ΔN (SBY24204), *pGAL-CSE4 MIF2-GFP* (SBY24212), *and pGAL-CSE4 mif2-*Δ*N-GFP* (SBY24216) yeast strains. Cells were arrested in G1 with alpha factor and released into galactose media to induce Cse4 overexpression and harvested after 120 minutes. The borders of the cells are outlined in white. The mCherry signal is weak in the merged images so the merges appear green. (C) Percentages of cells with more than two GFP foci in strains from panel (B) at different time points. At least 100 cells were imaged for each time point over three biological repeats. Error bars are standard deviation over three biological repeats. (D) Percentages of cells in anaphase determined by DAPI staining in strains from panel (A) and *pGAL-CSE4 mif2-*Δ*N mad3*Δ (SBY24224). At least 100 cells were imaged over three biological repeats for each time point. Error bars are standard deviation over three biological repeats. All error bars represent the standard deviation over three biological repeats. At least 3,000 DNA molecules were imaged for each time point from each biological replicate.

## Discussion

Here we report a time-series method for directly measuring the kinetics of kinetochore assembly at the single-molecule level and determining the order of subcomplex incorporation. Our method provides an unprecedented view of the kinetochore assembly process, revealing some surprising interdependencies between subcomplexes. The first two proteins to assemble were Ndc10, representing the CBF3 complex, and then the centromeric histone H3 variant Cse4, consistent with the role of CBF3 in Cse4 deposition (32,33,38). Next came one of the essential CCAN components, either Ame1 or Mif2 depending on whether Dsn1 was autoinhibited or not (as discussed below). Early assembly of these essential CCAN components was also expected given their specific affinity for centromeric nucleosomes (54,55). The remaining nonessential CCAN components followed, all with similar timing and efficiency, consistent with a highly interdigitated arrangement in the inner kinetochore (Figure 8A, top row) (16).

**Figure 8.**
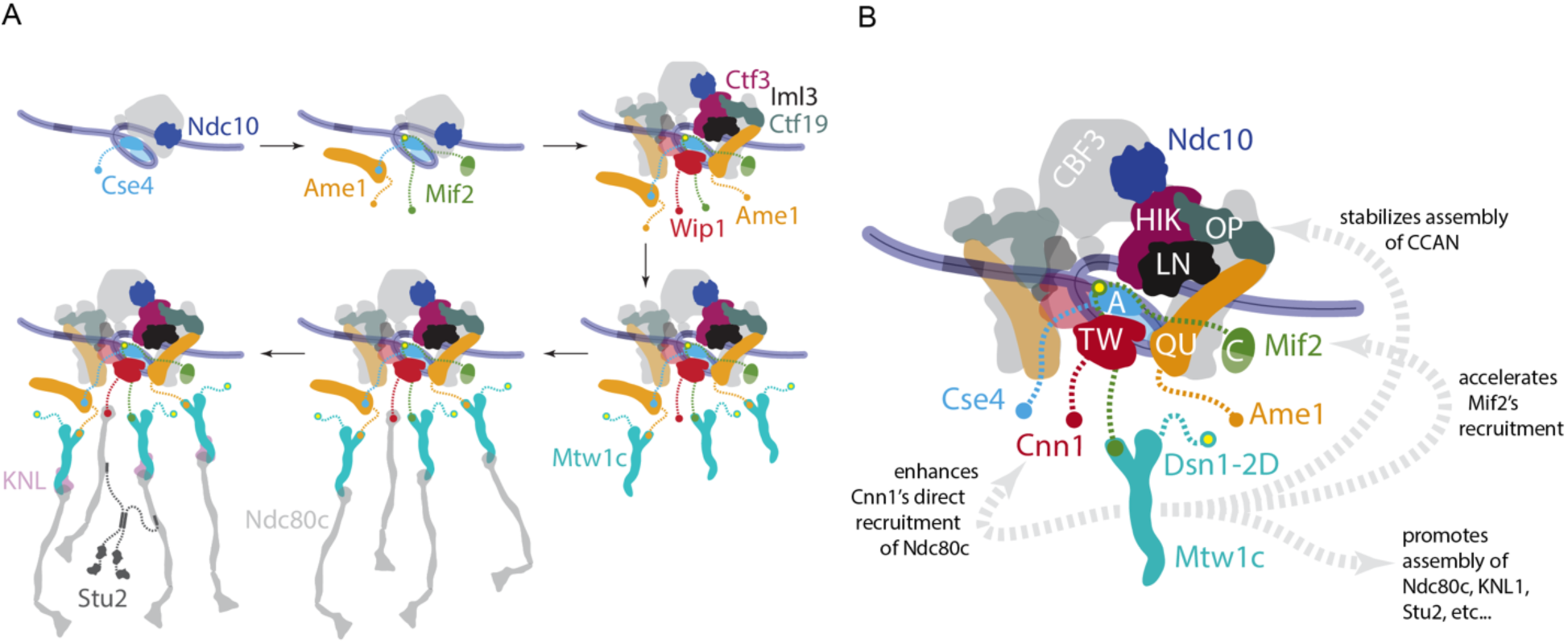
Summary of assembly order and interdependence between outer and inner kinetochore subcomplexes. (A) Schematic showing the order of assembly of indicated kinetochore components. *Top row:* Ndc10 and the centromeric histone H3 variant Cse4 assemble first, followed by the essential CCAN components, Ame1 and Mif2, and then the non-essential CCAN components. *Bottom row:* Among outer kinetochore components, Mtw1c assembles first, followed by Ndc80c, and then the KNL subcomplex and Stu2. (B) The phosphomimetic modification of Mtw1c promotes assembly in unexpected ways. Dsn1-2D enhanced direct recruitment of Ndc80c by Cnn1, increased the assembly efficiency of non-essential CCAN components, and accelerated Mif2 recruitment, in addition to its known role in promoting outer kinetochore assembly.

The timing of outer kinetochore assembly in our experiments was largely consistent with a tree-like hierarchical arrangement. Outer kinetochore subcomplexes were detectable after assembly in wild type lysates, but they assembled too inefficiently for kinetic analyses. We therefore used *dsn1-2D* to improve outer kinetochore assembly by relieving Mtw1c autoinhibition (24,30,45,56). The outer subcomplexes then assembled much more efficiently, as expected, after incorporation of the essential CCAN components. Mtw1c assembled first, followed by Ndc80c, then the KNL subcomplex and Stu2 (Figure 8A, bottom row).

Surprisingly, however, we found that the *dsn1-2D* modification of Mtw1c promotes assembly via at least three additional mechanisms not predicted by a simple tree-like hierarchy with independent branches (Figure 8B). First, *dsn1-2D* enhanced the contribution of Cnn1 to Ndc80c recruitment even though Cnn1 binds Ndc80c directly and has no known affinity for Mtw1c in budding yeast. Second, *dsn1-2D* increased the assembly efficiency of non-essential CCAN components (Ctf19, Ctf3, and Iml3), even though none of these directly bind the Mtw1 subcomplex. Third, *dsn1-2D* accelerated Mif2 recruitment, such that it preceded rather than followed Ame1, even though the Mtw1c-binding region of Mif2 is distant from the nucleosome-binding regions that anchor it to the inner kinetochore. Thus, assembly of the inner kinetochore can depend strongly on outer components, and outer kinetochore branches do not assemble entirely independently of one another. A key area of future work will be to uncover the biochemical and structural bases of these unexpected interdependencies.

Our data extend a recent discovery that Mif2 autoinhibition by its N-terminus can be relieved by binding to Cse4 (21) and suggest it can also be relieved by interacting with Mtw1. However, we did not detect Mtw1c arriving at the same time as Mif2 in the *dsn1-2D* lysates, suggesting that the autoinhibition is relieved in the nucleoplasm. It is therefore likely that the N-terminus of Mif2 exists in an equilibrium between being autoinhibited and being available to bind to Mtw1c. Once Mif2 autoinhibition is relieved, it stably associates with the centromere prior to stable Mtw1c incorporation. We also discovered that interaction between the Mif2 N-terminus and Mtw1c somehow improves recruitment of several CCAN subcomplexes (Iml3, Ctf3 and Wip1), even though they do not interact directly with either protein. This enhancement of CCAN recruitment, and the similar enhancement caused by *dsn1-2D* (described above) both indicate that kinetochore assembly is more interdependent than previously appreciated. Although the N-terminus of Mif2 is not essential, it becomes important for cell viability and kinetochore function when Cse4 is overexpressed. Although the precise underlying mechanisms remain to be uncovered, the sensitivity of *mif2-*Δ*N* strains to overexpression of Cse4 suggests that autoinhibition of Mif2 might contribute specificity to the kinetochore assembly process. Autoinhibition has also been demonstrated for several kinetochore components in addition to Mif2 and Mtw1c, including Ndc80c (57,58), CENP-E (59), and Spindly (60). Autoinhibition of numerous kinetochore subcomplexes, together with their highly interdependent recruitment, might prevent premature assembly of functional kinetochores at ectopic chromatin sites or in soluble forms, where they could interfere with accurate chromosome segregation. We hope the methods developed here will ultimately help test this idea and uncover more principles underlying the hierarchical assembly of kinetochores specifically at centromeres, using yeast and potentially mammalian cell extracts.

## Data Availability

The data underlying this article will be shared on request to the corresponding author.

## Supplementary Data statement

Supplementary Data are available at NAR Online

## Acknowledgements

We are grateful to the Biggins and Asbury lab members for the helpful suggestions for this study and their comments on the manuscript. We also thank the Cellular Imaging team, especially Lena Schroeder and Hoku West-Foyle, for all their help.

## Author Contributions

Conceptualization (C.Hu, C. Asbury, and S. Biggins), data curation (C.Hu, A. Andrade Latino, S. Biggins), formal analysis (C.Hu, C. Asbury, S. Biggins), funding acquisition (C. Hu, C. Asbury, S. Biggins), investigation (C.Hu, A. Popchock, and S. Biggins), project administration (S. Biggins), resources (C.Hu, A. Andrade Latino, S. Biggins), software (C.Hu, A. Popchock, and C. Asbury), supervision (S. Biggins), validation (C.Hu and S. Biggins), visualization (C.Hu, C. Asbury, and S. Biggins), writing–original draft (C.Hu and S. Biggins), writing–review and editing (C.Hu, C. Asbury, S. Biggins).

## Funding

P30 CA015704/CA/NCI NIH HHS/United States

NIH R35GM134842/HHS | National Institutes of Health (NIH) to C.A.

R35 GM149357/GM/NIGMS NIH HHS/United States to S.B.

A.R.P. was supported by postdoctoral fellowship NIH F32GM136010

HHMI/Jane Coffin Childs Memorial Fund postdoctoral fellowship to C.H.

HHMI to S.B.

## Conflict of Interest Disclosure

The authors declare no conflict of interest.

## Notes

### Competing Interest Statement

The authors have declared no competing interest.

