## Supplemental Figures for "Direct observation of interdependent and hierarchical kinetochore assembly on individual centromeres"

Hu et al., Fig. S1

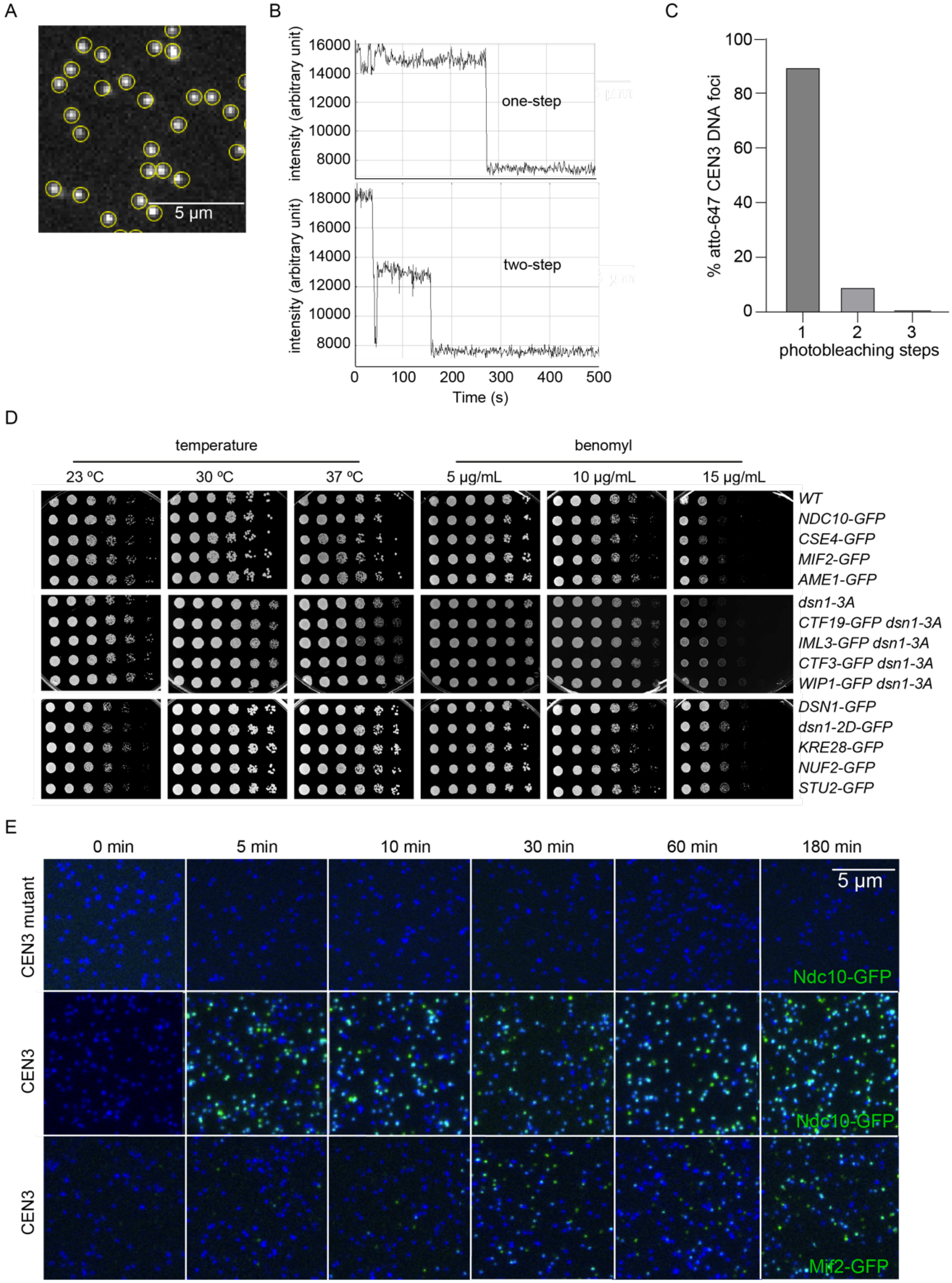

**Supplementary Figure S1**. **GFP-tagged kinetochore proteins bind specifically to CEN3 DNAs.**

(A) Representative image of CEN3 DNAs. Yellow circles indicate the automatic selection of each DNA focus detected by fluorescence (Atto-647).
(B) Representative Atto-647 fluorescence intensity trace over time showing one-step (top) and two-step photobleaching (bottom).
(C) Quantification of photobleaching steps for each DNA focus from panel (A) analyzed in MATLAB using previously a described step-finding algorithm (27).
(D) Five-fold serial dilutions of yeast strains containing the indicated GFP tags. Strains include NDC10-GFP (SBY22903),CSE4-GFP (SBY22195), MIF2-GFP (SBY22094), AME1-GFP (SBY22119), dsn1-3A (SBY14171), CTF19-GFP dsn1-3A (SBY24416), IML3-GFP dsn1-3A (SBY24442), CTF3 dsn1-3A (SBY24432), WIP1-GFP dsn1-3A (SBY24440), DSN1-GFP (SBY22153), dsn1-2D-GFP (SBY22159), KRE28-GFP (SBY24188), NUF2-GFP (SBY23256), STU2-GFP (SBY22135), and the untagged parental control (wild type, SBY4). Cells were plated on YPD or benomyl media. Cells were grown on YPD at the indicated temperatures for 48 hours or at the indicated benomyl concentrations at 23 ºC.
(E) Representative TIRFM images of Mif2-GFP (SBY22094) and Ndc10-GFP (SBY22903) (green foci) after a time series incubation with CEN3 DNAs (blue foci, middle and bottom rows) and CEN3 mutant DNAs (blue foci, top row). Yeast were arrested in mitosis using benomyl prior to harvesting. The lysates were washed off before imaging.

Hu et al., Fig. S2

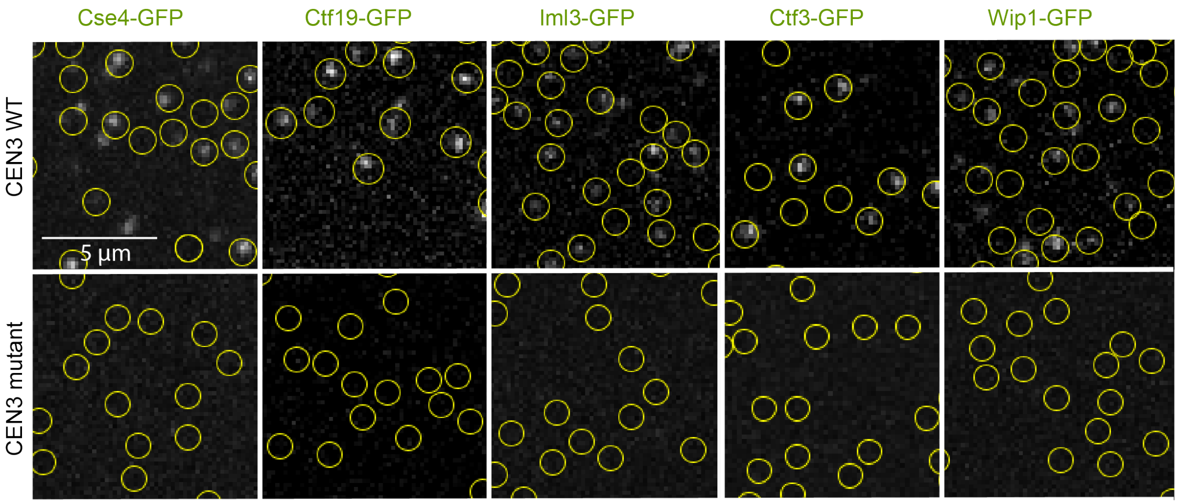


**Supplementary Figure S2**. **GFP tagged kinetochore proteins bind specifically to CEN3 DNAs.**

Representative images of CEN3 or CEN3 mutant DNAs (at locations indicated by yellow circles) after kinetochore assembly in lysates made from strains with GFP-tagged Cse4 (SBY22195), Ctf19 (SBY22116), Iml3 (SBY22199), Ctf3 (SBY22203), and Wip1 (SBY22207). Yeast cells were arrested in mitosis with benomyl prior to harvesting the lysates. The lysates were washed off the slides after 180 minutes of incubation at room temperature (21-23 ºC) before imaging.

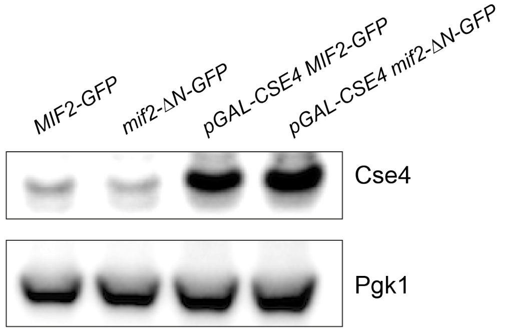


**Supplementary Figure S3**. **Cse4 levels are similar in *MIF2* and *mif2-ΔN* strains.**

Yeast lysates were prepared from MIF2-GFP (SBY22095), mif2-ΔN (SBY23249), pGAL-CSE4 MIF2-GFP (SBY24220), and pGAL-CSE4 mif2-ΔN-GFP (SBY24222) yeast strains grown in galactose. Immunoblotting with anti-Cse4 antibodies indicates that the levels of wild type and overexpressed Cse4 are similar in MIF2 and mif2-ΔN strains.
